# Data and model considerations for estimating time-varying functional connectivity in fMRI

**DOI:** 10.1101/2021.07.28.454017

**Authors:** C Ahrends, A Stevner, U Pervaiz, ML Kringelbach, P Vuust, M Woolrich, D Vidaurre

## Abstract

Functional connectivity (FC) in the brain has been shown to exhibit subtle but reliable modulations within a session. One way of estimating time-varying FC is by using state-based models that describe fMRI time series as temporal sequences of states, each with an associated, characteristic pattern of FC. However, the estimation of these models from data sometimes fails to capture changes in a meaningful way, such that the model estimation assigns entire sessions (or the largest part of them) to a single state, therefore failing to capture within-session state modulations effectively; we refer to this phenomenon as the model becoming static, or model stasis. Here, we aim to quantify how the nature of the data and the choice of model parameters affect the model’s ability to detect temporal changes in FC using both simulated fMRI time courses and resting state fMRI data. We show that large between-subject FC differences can overwhelm subtler within-session modulations, causing the model to become static. Further, the choice of parcellation can also affect the model’s ability to detect temporal changes. We finally show that the model often becomes static when the number of free parameters that need to be estimated is high and the number of observations available for this estimation is low in comparison. Based on these findings, we derive a set of practical recommendations for time-varying FC studies, in terms of preprocessing, parcellation and complexity of the model.

**Highlights:**

- Time-varying FC models sometimes fail to detect temporal changes in fMRI data
- Between- and within-subject FC variability affect model stasis
- The choice of parcellation affects model stasis in real fMRI data
- The number of observations and free parameters critically affect model stasis Keywords: fMRI; time-varying FC; Hidden Markov Model (HMM); resting state

## 1 Introduction

Neural circuits across multiple brain areas integrate into large-scale brain networks in order to accomplish complex cognitive functions. Just like smaller populations of neurons underlying these networks flexibly synchronise and desynchronise their oscillatory firing patters to communicate (Fries, 2005), large-scale brain networks must also be able to fluctuate dynamically and change over time (Breakspear, 2017; Calhoun *et al*., 2014), enabling flexible neuronal communication and functioning across the entire brain. Arguably, this is reflected in the data as some form of synchrony in the activity across areas, which is typically referred to as functional connectivity (FC). In fMRI, FC can be derived by measuring how different areas coactivate in their blood oxygen level dependent (BOLD) signal. Understanding these temporal changes in FC (i.e. time-varying FC) in fMRI can help to address a range of questions, from the theoretical study of human cognition to a better characterisation of different neurological and psychiatric diseases.

There are several approaches to modelling time-varying FC in fMRI; for a recent review, see Lurie *et al*. (2019). One avenue is the use of state-based models that estimate time-varying FC as a temporal sequence of brain “states”. However, in fMRI these models are not always effective to detect changes in FC over time. Sometimes, the estimation leads to entire sessions collapsing into one single state, with no changes within session —so that the model becomes static; that is, all the explanatory power of the model is focussed on explaining differences between subjects or sessions, instead of within-session modulations. The reason behind this behaviour is an open question: it could be because there are no temporal changes in the data; or it could be because, even if there are temporal changes, the estimation is unable to detect them. While some studies have claimed that there is insufficient evidence that BOLD FC is dynamic (Hindriks *et al*., 2016; Liégeois *et al*., 2017; Lindquist *et al*., 2014), several studies have shown that dynamic aspects of FC are relevant for behaviour (Cabral *et al*., 2017; Fornito and Bullmore, 2010; Gonzalez-Castillo and Bandettini, 2018; Liegeois *et al*., 2019; Vidaurre *et al*., 2017; Voytek and Knight, 2015; Xie *et al*., 2018) and that they can add important information not contained in time-averaged FC (Vidaurre *et al*., 2021). These findings suggest that temporal variation is present in BOLD FC and that it carries meaningful information. We here address the question of how to quantify this variability effectively, and under which conditions the models become static (i.e. when “model stasis” occurs), since a deeper understanding of the issue can help us to configure these models to work more optimally.

Assuming temporal FC changes exist in the data, why would a time-varying FC model fail to detect them? One possible explanation is in the nature of the data. We here refer to factors that affect the data as the **data hypothesis**. In particular, since unsupervised, data-driven time-varying FC models aim at describing the most salient patterns in the data, within-session fluctuations might just be too subtle, with overall differences between subjects being more dominant (Lehmann *et al*., 2017). That is, if *between*-subjects differences are larger than *within*-session FC modulations (i.e. changes over time within a subject’s scanning session), a data-driven model will naturally prefer to focus on the between-subjects variability instead of the temporal variability. As we will show, the balance between these two aspects of variability (between-subjects and within-session) depends on the preprocessing pipeline, in particular on the choice of a parcellation (Eickhoff *et al*., 2018; Pervaiz *et al*., 2020; Popovych *et al*., 2021), and how fine-grained it is. Another explanation relates to challenges in estimating the model; i.e. if the model inference has problems in finding within-session modulations. We refer to this explanation as the **estimation hypothesis**, which, in particular, might occur when the number of free parameters to estimate in the model is too large in comparison to the available number of volumes or time points (across subjects).

In the present study, we simulated data with varying amounts of variability between and within subjects, and we fitted models to a real dataset in different parcellations. We hypothesise that large between-subject variability and small within-subject (temporal) variability cause the time-varying FC model to become static and that this effect depends on the parcellation (data hypothesis). We further hypothesise that fewer observations and more free parameters, in fact a small ratio of number of observations to free parameters, cause the time-varying FC model to become static (estimation hypothesis). We finally provide some recommendations for the estimation of time-varying FC based on these points.

## 2 Material and Methods

### 2.1 Data and parameters

#### 2.1.1 HCP dataset and preprocessing

We used resting state EPI scans of the first 200 participants from the Human Connectome Project S1200 (HCP, (Smith *et al*., 2013b; Van Essen *et al*., 2013)), an open-access dataset of MRI data. Time-varying FC has previously been demonstrated in this dataset using a wide array of different approaches (Battaglia *et al*., 2020; Casorso *et al*., 2019; Choe *et al*., 2017; Dai *et al*., 2019; Liegeois *et al*., 2019; Riccelli *et al*., 2017; Sporns *et al*., 2021; Vidaurre *et al*., 2017; Zalesky *et al*., 2014; Zamani Esfahlani *et al*., 2020), making it a suitable example to evaluate model stasis. The dataset consists of structural and functional MRI data of 1200 healthy, young adults (age 22-35). Each participant completed four resting state scans. We here only used data from the first resting state scanning session of each participant. Data were acquired as described in the HCP public protocols, which can be found in Van Essen *et al*. (2012). Briefly, scans were acquired in a 3T MRI scanner, using multiband echo planar imaging sequences with an acceleration factor of 8 at 0.73 seconds repetition time (TR) and a spatial resolution of 2×2×2 mm for functional scans. Resting state scans lasted 15 minutes.

Data were preprocessed following the HCP preprocessing pipelines for resting-state fMRI (Glasser *et al*., 2013; Smith *et al*., 2013a). In brief, after “minimal” spatial preprocessing and surface projection to transform data into grayordinate space, the data were temporally preprocessed using single-session Independent Component Analysis (ICA, using FSL’s MELODIC; (Beckmann, 2012)), as well as classification and removal of noise components using FSL’s FIX (Griffanti *et al*., 2014; Salimi-Khorshidi *et al*., 2014).

#### 2.1.2 Parcellations and time course extraction

Group ICA parcellations estimate a data-driven functional parcellation on the group level, which are subsequently regressed onto each subject’s individual functional scans to obtain subject-specific versions of group ICs and their time courses. Group ICA parcellations were created for a varying number of parcels (we here used the variants created for 50 and 100 parcels, GroupICA50 and GroupICA100) using multi-session spatial ICA on the temporally concatenated data. The time series for each participant were extracted using dual regression (Beckmann *et al*., 2009). The Group ICA parcellations and corresponding time series are publicly available from the HCP repository (https://db.humanconnectome.org).

PROFUMO (Harrison *et al*., 2015) is a similar approach to Group ICA, but it estimates group-as well as subject-level maps simultaneously, allowing it to better capture individual variability in FC (Bijsterbosch *et al*., 2018). In PROFUMO, between-subject differences in (time-averaged) FC are therefore expected to be higher compared to the group ICA approach. We used a PROFUMO parcellation of 50 parcels, PROFUMO50.

As *a priori* defined functional parcellation, we used the Yeo parcellation (Schaefer *et al*., 2018). This parcellation was created using a gradient-weighted Markov Random Field on a separate dataset of resting-state fMRI recordings of 1489 participants. This approach produces parcels which are similar in terms of function and connectivity. We here used the grayordinate version of this parcellation consisting of 100 parcels (Yeo100 parcellation).

As an anatomical parcellation, we used the Desikan-Killiany atlas (Desikan *et al*., 2006). This atlas originally consists of 62 anatomically delineated cortical regions. The atlas was projected into grayordinate space and 18 subcortical regions were added, as described in Deco *et al*. (2021). This resulted in 80 parcels (DK80 parcellation). Time courses in this parcellation were extracted as the mean across grayordinates belonging to each parcel.

Beside runs that use the full parcellations, we also ran the models on subsets of each parcellation to vary the number of free parameters in the model (as described under Section 2.3). In these reduced runs, we randomly chose a subset of 10, 25, or 50 parcels from a parcellation, which time series were subsequently fed to the model. As an alternative strategy to reduce the number of free parameters in the model, we also tested the effects of reducing the original data dimensionality using Principal Component Analysis (PCA) or by modelling each HMM-state as probabilistic PCA model (“HMM-PCA”) (Vidaurre, 2021).

To mimic properties of more ordinary datasets, we also varied the number of subjects (*S*) between 50, 100, and 200, the number of time points (*T*) per subject between 200, 500, and 1200 time points, and the fraction *R* of the sampling rate at 1.37 Hz (original rate *R* = 1, equivalent to TR of 0.73 s), 0.68 Hz (half of the original rate 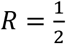, equivalent to TR of 1.46 s), and 0.46 Hz (one third of the original rate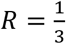, equivalent to TR of 2.19 s). The number of observations *O* used in the model is the total amount of time points: *O* = *S* * *T* * *R*. In our analysis, we modelled only the effect of the number of observations *O*, rather than the effects of the number of subjects, of time points, and of the sampling rate separately.

Time course extraction results in one matrix of dimensions *T* x *N* per subject, where *N* is the number of parcels. To compute time-varying FC, we concatenated the time series across subjects, resulting in a matrix of (*S* x *T*) x *N*. The input and size at this step varies with the parcellation, i.e. *N* = 50 for the GroupICA50 and PROFUMO50 parcellations, *N* = 80 for the anatomical DK80 parcellation, and *N* = 100 for the GroupICA100 and Yeo100 parcellations in the full runs (i.e. with all regions or components). In the reduced runs, *N* corresponds to the number of randomly chosen parcels from each parcellation (10, 25, or 50 parcels). We then standardised these time series row-wise by rescaling them so that the time course of each parcel has a mean of 0 and a standard deviation of 1.

#### 2.1.3 Simulations

To be able to test the different levels of between-subject and within-session variability, we simulated new datasets based on the HCP data, where we introduced differing amounts of between-subject and within-session variability into the generating model. This was done by generating new time series from a combination of synthetic covariance matrices, representing either time-invariant (subject-specific) FC matrices or time-varying FC matrices that activate or deactivate at different time points:

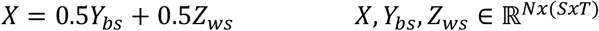

Here, *X* is the synthetic time series containing variability both between subjects and within sessions, *Y*_*bs*_ is the synthetic time series containing only variability between subjects, and *Z*_*ws*_ is the synthetic time series containing only variability within sessions. In this notation, *X, Y*_*bs*_, and *Z*_*ws*_ all represent subjects’ individual time series that have been concatenated.

The time series *Y*_*bs*_, containing only variability between subjects, was generated by randomly sampling from a Gaussian distribution with mean 0 and a different synthetic covariance matrix per subject:

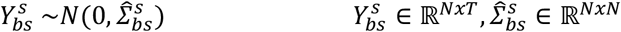

where 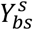 is the time series for subject *s*, and 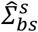is the (symmetric, positive-definite) covariance matrix of subject *s*, i.e. containing FC information specific for this subject and different from the others.

The time series *Z*_*ws*_, with only variability within a session, was obtained by sampling from an HMM distribution:

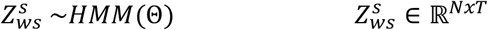

where 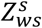is the time series for subject *s* . Critically, 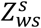contains only within-session variability, since the HMM parameters Θ are at the group level (i.e. equal for all subjects). More specifically, when a given state *k* is active, 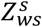is sampled from a Gaussian distribution with mean 0 and a state-specific synthetic covariance 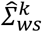:

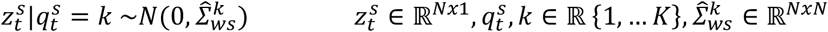

Therefore, 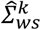 accounts for state-specific variability, which depends on the currently active state 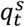. The currently active state 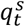 depends on which state was active at the previous time point 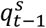 . The states are sampled from a categorical distribution with the parameters A, which are the transition probabilities of the HMM (A_*k*_ indicating the k -th row of the transition probability matrix):

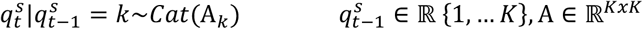

To create the covariance matrices 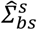 and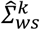, we first decomposed the real covariance matrix of the first subject of the HCP dataset into its singular values:

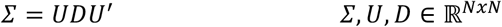

where *Σ* is the covariance matrix of the first subject of the real dataset (HCP resting-state fMRI dataset) in GroupICA50 parcellation, *U* are the singular vectors of the covariance matrix and *D* contains the singular values of the covariance matrix.

We created synthetic covariance matrices 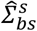 for all subjects *S* by multiplying the original singular values *D* with subject-specific singular vectors 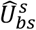, which we created by randomly perturbing *U*.

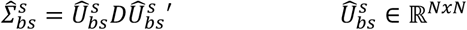

Similarly, to create the covariance matrices 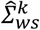 for all states *K*, we multiplied the original singular values *D* with state-specific singular vectors 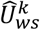:

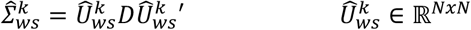

For each subject *s*, the noisy singular vectors 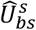 were generated by multiplying the original singular vectors *U* element-wise with a subject-specific Gaussian noise matrix *ψ*^*s*^ and adding this product to the original vectors *U*:

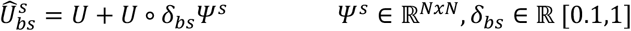

The Gaussian noise matrix *ψ*^*s*^ is scaled by the parameter *δ*_*bs*_, which defines the final amount of between-subject variability contained in the synthetic time series *Y*_*bs*_. Similarly, for each state *k*, we generated the noisy singular vectors 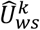 by multiplying the original singular vector *U* element-wise with a state-specific Gaussian noise matrix *ψ*^*k*^:

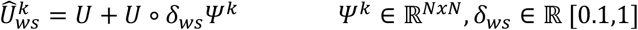

This Gaussian noise matrix *ψ*^*k*^ is scaled by the parameter *δ*_*ws*_, which defines the amount of within-session variability contained in the synthetic time series *Z*_*ws*_.

We varied the parameters *δ*_*bs*_ and *δ*_*ws*_ between 0.1 and 1 in steps of 0.1. A small value for *δ* _*b*_s results in a time series *Y*_*bs*_, in which all subjects’ time-averaged FC are similar. A large value for *δ*_*bs*_, on the other hand, results in a time series *Y*_*bs*_, in which subjects’ FC matrices are very different from each other. A small value for *δ*_*ws*_ results in a time series *Z*_*ws*_, in which FC almost does not vary over time (i.e. FC is essentially static). A large value for *δ*_*ws*_, on the other hand, results in a time series *Z*_*ws*_, in which FC varies greatly over time.

We generated time series from all combinations *X* of *Y*_*bs*_ and *Z*_*ws*_, resulting in 100 simulated time series. We then used these time series as input to compute time-averaged FC, as described under 2.2, and to the time-varying FC model to evaluate the model’s stasis, as described under 2.3.

### 2.2 Time-averaged functional connectivity and FC similarity

To compute time-averaged functional connectivity, Pearson’s correlation was computed for each pair of regions (Smith *et al*., 2013c). The resulting *N* x *N* matrices represent the time-averaged FC of each scanning session within each parcellation. In order to assess how consistent these FC networks were for each of the parcellations, we estimated the network similarity across scanning sessions. This was done by first calculating the group average of the time-averaged FC in each parcellation, then unwrapping the upper triangular elements of this group average FC matrix into a 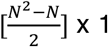 x 1 vector, and correlating this group-level vector with the corresponding vectors of the session-specific FC matrices. For each parcellation, FC similarity was thus defined as the correlation between the group mean FC, and the FC of all individual scanning sessions.

### 2.3 Time-varying functional connectivity: Hidden Markov Model (HMM) and model stasis

We used the Hidden Markov Model (HMM; Vidaurre *et al*. (2016); Vidaurre *et al*. (2017)) to describe time-varying FC. The HMM is a type of state-based model that estimates a sequence of states and a probability distribution for each state, such that each time point in the time series is assumed to have been generated from its assigned state distribution. The HMM has been used to estimate time-varying FC on fMRI and MEG data in previous work (Quinn *et al*., 2018; Stevner *et al*., 2019; Vidaurre *et al*., 2016; Vidaurre *et al*., 2017).

We used a version of the HMM that assumes a multivariate Gaussian distribution per state, with *K* = 6 states for the simulated data and *K* = 12 for the HCP data. In order to focus on FC, each state was here defined in terms of its covariance only (Vidaurre, 2021), i.e. without explicitly modelling the mean (or amplitude). Once the model was estimated, we computed the fractional occupancy (FO), defined as the proportion that each state occupies in the time series of a particular subject. We used FO as indicator of the model becoming static. This is illustrated in **Figure** 1B. In the example, different states are assigned to portions of the time series of Subject 3, resulting in a small FO percentage for each of the states. For Subject 4, however, a single state (#1) is assigned to all time points of this subject, i.e. the FO of state #1 in Subject 4 is 100%. This means that the model effectively fails in finding any temporal changes in functional connectivity for this subject, describing only the more salient difference between all time points of this subject compared to the other subjects. To evaluate the model’s overall stasis, we then used the maximum FO value of each subject and computed the average across the group. In practice, this means that a model that assigns states only to entire subjects, such as in the example with Subject 4, will have a mean maxFO of 100%. On the other hand, a model that finds recurring states over time that are perfectly equally distributed across time points of all subjects will have a mean maxFO of approximately 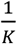 (i.e. each of the *K* states occupies on average the same amount of each subject’s time series). We here used stasis, as measured by the model’s mean maxFO, as an indicator of how well a time-varying FC model is able to estimate temporally recurring states.

**Figure 1.**
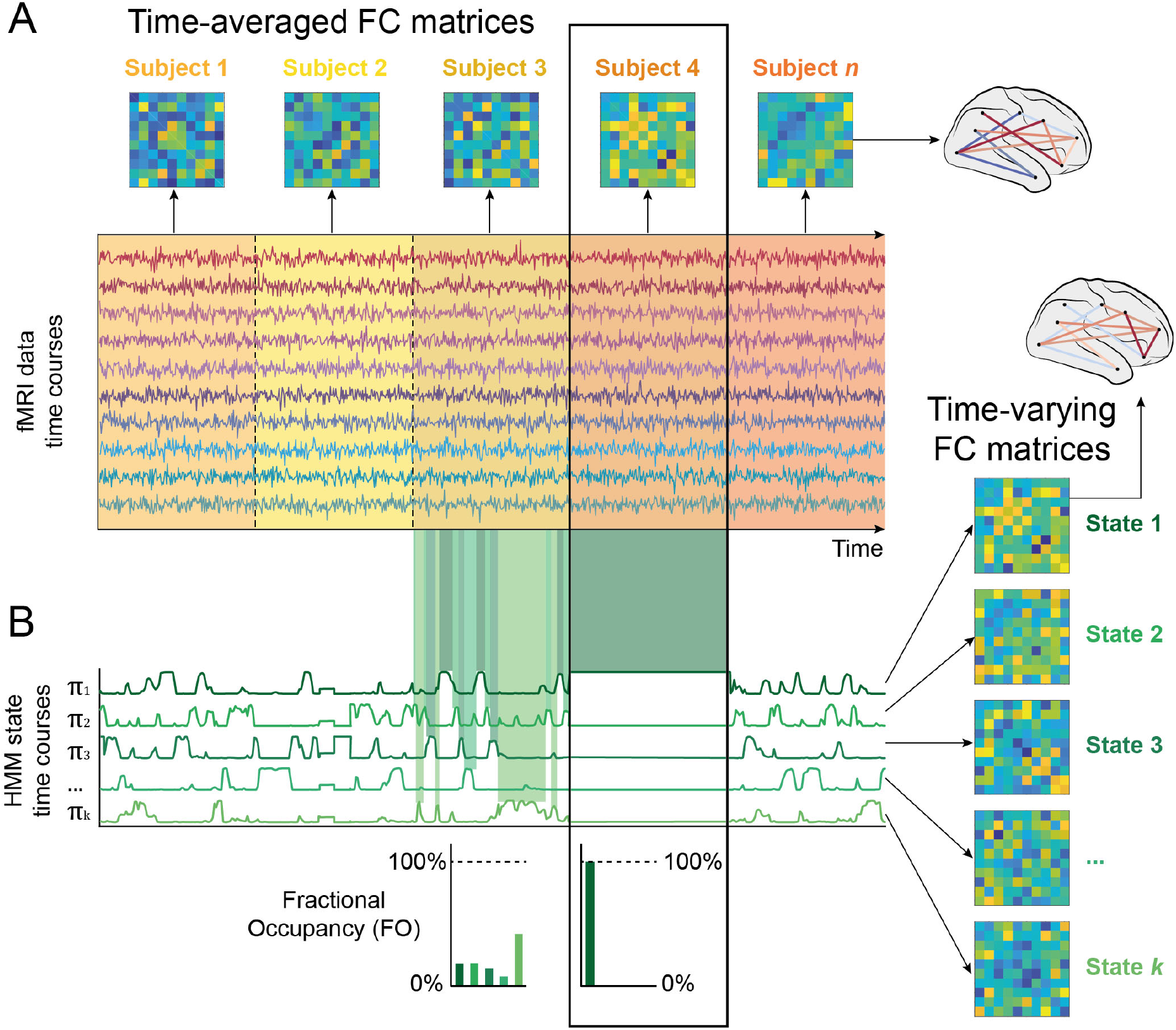
Between-subject variability in time-averaged FC may affect stasis in a time-varying FC model. **A)** Time-averaged FC (regions by regions) matrices for each subject were obtained by pairwise correlating time courses of all parcels from each subject. Subjects are represented in the time series as different colours. As observed, the time-averaged FC matrix from Subject 4 is very different from the time-averaged FC matrices of the other subjects. **B)** Given a prespecified number of states K, the Hidden Markov Model (HMM) estimates both the state-specific FC matrices and when the states become active. In the example for Subject 3, all states transiently occur and recur over time. In opposition to this temporal recurrence in Subject 3, the HMM time course for the time points corresponding to Subject 4 stays stable at a high probability for state 1. The temporal recurrence of states can be measured by their fractional occupancy (FO), indicating the proportion of the entire time series that a given state occupies. In this example, state FOs for Subject 3 indicate that all states take up a similar amount of the time series with certain states being relatively more prevalent than others. In Subject 4, however, the FO of state 1 is at 100% while all others are at 0%, since state 1 occupies the entire time series of this subject. This is summarised by the term “stasis”: The model is static when one state’s FO approaches 100% and all others are close to 0%.

To test our estimation hypothesis, we calculated the number of free parameters of each model. If this number is too large in comparison to the number of observations *O*, the estimation may become statistically challenging. The number of free parameters *DF* in a HMM with *K* states, each defined by a full covariance matrix but without modelling the mean, and *N* parcels can be computed as

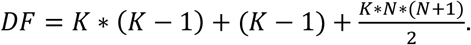

We implemented the model using the HMM-MAR toolbox available at https://github.com/OHBA-analysis/HMM-MAR in MATLAB (Mathworks, 2016). Although the HMM is only one example of a time-varying FC model, the concepts discussed here are likely to apply also to other models of time-varying FC.

### 2.4 Structural Equation Modelling (SEM)

To provide a synthesis of the hypothesised relationships, we modelled all effects in a structural equation model (SEM). SEM characterises the causal links between variables, which are combined in a network of structural equations. In these structural equations, the relationships between variables are explicitly declared. Each variable can be declared to have a direct effect on an outcome variable or an indirect effect by declaring this variable simultaneously as an outcome and a predictor variable. A single variable can also have both a direct and an indirect effect on an outcome variable. Here, we combined a series of linear models and linear mixed effects models in a piecewise SEM, also called confirmatory path analysis (Shipley, 2000). Rather than estimating coefficients in a single variance-covariance matrix as in traditional SEM, piecewise SEM first estimates each part of the model independently before evaluating them at the level of the full model. This allows increased flexibility on the level of the constituting parts of the SEM in terms of their distributions, making it possible e.g. to include random effects in parts of the model.

We fitted two separate SEMs: one to the outcomes of HMMs run on simulated data and one to the outcomes of HMMs run on the real (HCP) data. In both SEMs, there are two parts. The first part constitutes the effect of the observed variables on FC similarity, and the second part links the observed variables and FC similarity to mean maxFO as an indicator of the HMM’s model stasis. In the SEM on simulated data, the first part modelled the effects on FC similarity of the number of observations *O* (which here depends only on the number of subjects *S*) and of between-subject variability (the value of the parameter *δ*_*bs*_, described under 2.1.3). The second part modelled the effects on model stasis of the number of observations *O*, of FC similarity, of within-session variability (the value of the parameter *δ*_*ws*_, described under 2.1.3) and of the inverse of the number of free parameters *DF* (which varies based on the number of parcels *N* from each parcellation). The number of observations *O* has therefore both a direct effect on model stasis and an indirect effect via FC similarity. In the SEM on real data, the first part modelled the effect on FC similarity of the number of observations *O* (which here varies based on the number of subjects *S*, the number of time points *T*, and the sampling rate *R*). We additionally included a random intercept for the different parcellations in this model. In the second part, we modelled the effects on model stasis of the number of observations *O*, of the inverse of the number of free parameters *DF* (which varies based on the number of parcels *N* from each parcellation), of their interaction, and of FC similarity. In this SEM, we included both a random intercept and random slope of the effect of FC similarity for each parcellation. The number of observations *O* has again both a direct and an indirect (via FC similarity) effect on model stasis.

We used the piecewiseSEM-package (Lefcheck, 2016) in R (R Core Team, 2020) to fit the SEM models as a combination of linear and linear mixed effects models.

## 3 Results

We address the factors from the two hypotheses (3.1 Data hypothesis, **Error! Reference source not found**. Estimation hypothesis) one by one, distinguishing between results from simulated data and real data (HCP data). Statistics from the full structural equation models (SEM) are summarised under 3.3.

### 3.1 Data hypothesis

We first investigated which aspects of the data influence the ability of a time-varying FC model to detect temporal changes in FC (data hypothesis). Namely, we tested the effects of between-subject variability and of within-session variability on FC similarity and on model stasis in simulated time series (3.1.1). We then focussed on the effect of the parcellation used to extract time series from the HCP resting state data on FC similarity, on model stasis, and on the relationship between them (3.1.2).

#### 3.1.1 Between-subject and within-session variability in simulated time series affect model stasis

Here, we show on synthetic data that large differences between subjects or small differences over time can cause the time-varying FC model to become static.

In order to address the question of variability in the data, we simulated new data with different degrees of between-subject and within-session variability (described under 2.1.3). First, we calculated FC similarity of these new FC matrices, which confirmed that this measure robustly reflects between-subject variability *δ*_*bs*_, independently of within-session variability *δ*_*ws*_ (see **Figure 2**A, top panel). In the full structural equation model (SEM), FC similarity is near perfectly explained (standardised coefficient of -0.97, *p*<0.0001) by between-subject variability *δ*_*bs*_ . We can therefore assume that, in the real data, FC similarity is a reliable proxy for between-subject variability.

**Figure 2.**
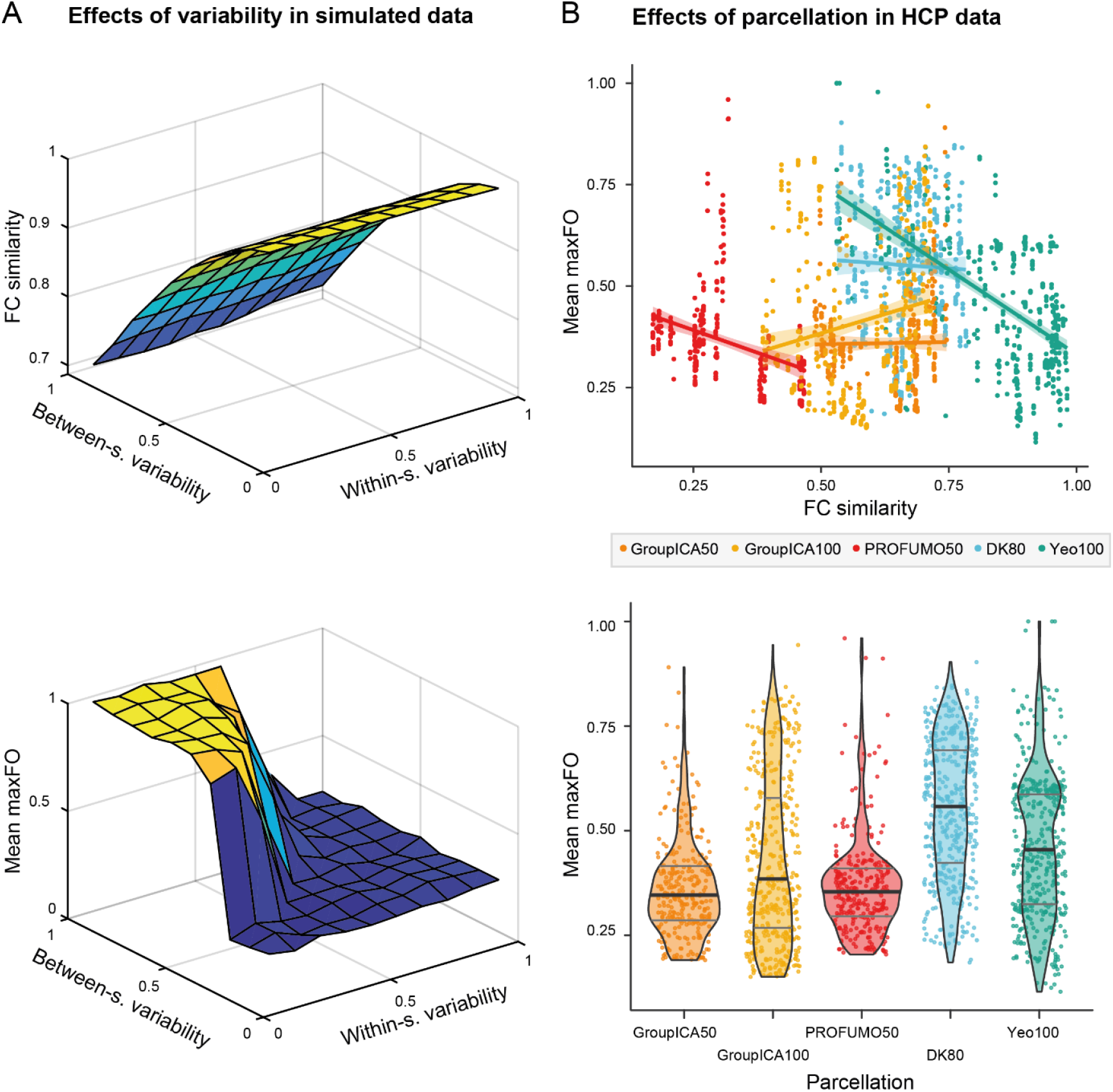
Evidence for the data hypothesis. **A)** In the simulated data, between-subject variability but not within-session variability affects FC similarity between subjects (top panel). The bottom panel shows how between-subject and within-session variability affect model stasis (as measured by mean maxFO) in a time-varying FC model. In the graph area where between-subject noise is high and within-subject noise is low, the model is static (yellow area). **B)** In the real data, FC similarity and model stasis depend on the parcellation. We here represent each parcellation with a different colour. In the top panel, we illustrate the linear regression line and corresponding 95% confidence interval between FC similarity and model stasis (represented by the mean maxFO statistic) within each of the parcellations. The graph shows how both the position and the slopes for these regression lines are different between parcellations. In the bottom panel, we show the distribution of mean maxFO values within each of the parcellations. The thick black line within each violin plot indicates the mean value of mean maxFO in the respective parcellation and the grey lines indicate their interquartile range. Dots within each parcellation correspond to runs with different dimensionality parameters as described under 2.1, i.e. different numbers of subjects S, time points T, sampling rates R, and (subsets of) parcels N.

FC similarity was not significantly affected by the number of observations *O* (i.e. by varying the number of subjects *S*) (coefficient: -0.02, *p*=0.99). As hypothesised, model stasis depends on both between-subject and within-session variability, where high between-subject and low within-session variability cause the model to become static. Decreasing differences between subjects and increasing temporal variability in the data lead to a lower rate of model stasis. This is shown for an exemplary solution in **Figure 2A**, bottom panel. In the full model, the effects of between-subject and within-session variability are of a similar magnitude, with standardised coefficients of -0.53 (*p*<0.0001) for FC similarity and -0.54 (*p*<0.0001) for within-session variability.

In summary, this indicates that the between-subject vs. within-session variability balance is an important contributor to model stasis. That is, if subjects in the dataset are very dissimilar, differences across time points need to be large in order for a time-varying FC model to be able to identify dynamically changing states. In real datasets, it may therefore be important to work towards high similarity between subjects while retaining temporal variation as much as possible during preprocessing. One central factor in achieving this may be the choice of parcellation, which we tested next (3.1.2).

#### 3.1.2 The parcellation affects FC similarity, model stasis, and the relationship between them

We next investigated the effect of the parcellation on FC similarity, on model stasis, and on the relationship between them. As we will see, FC similarity does not simply explain model stasis, but the choice of parcellation can strongly affect FC similarity, model stasis, and the relationship between these two variables.

Time courses were extracted from the HCP data in five different parcellations: We used three data-driven functional parcellations (GroupICA50, GroupICA100, and PROFUMO50 (Beckmann *et al*., 2009; Harrison *et al*., 2015)), one *a priori* defined functional parcellation (Yeo100 (Schaefer *et al*., 2018)), and one anatomical parcellation (DK80 (Deco *et al*., 2021; Desikan *et al*., 2006)). As shown in **Figure 2**B, the choice of parcellation affects FC similarity, model stasis (as measured by mean maxFO), and the relationship between them. We included the parcellation as random intercept in the first part of the full SEM (predicting FC similarity) and as random intercept and slope in the second part of the full SEM (predicting model stasis). This increased the variance explained by the full SEM as compared to a model excluding the effect of parcellation by 32% (*R reduced 2* =0.40, *R full2* =0.72). In the full SEM, the remaining effect of FC similarity on model stasis, i.e. the fixed effect not depending on parcellation, is not significant (coefficient -0.06, *p*=0.80). This indicates that the effect of FC similarity on model stasis is not as straightforward as we hypothesised, but strongly depends on the parcellation. The parcellations that, on average, created the most similar time-averaged FC matrices between subjects, increased model stasis the most.

Besides the parcellation, FC similarity is also significantly explained by the number of observations *O*, yielding a coefficient of 0.23 (*p*<0.0001). Across all runs, the parcellations ranked from least to most model stasis are: 1. GroupICA50 (M: 0.36 ± 0.12 S.D.), 2. PROFUMO50 (M: 0.37 ± 0.12 S.D.), 3. GroupICA100 (M: 0.41 ± 0.20 S.D.), 4. Yeo100 (M: 0.46 ± 0.17 S.D.), 5. DK80 (M: 0.55 ± 0.16 S.D.). On average, the three data-driven functional parcellations used here outperformed both the example of a functional and the example of an anatomical parcellation, in the model’s ability to detect dynamic changes in FC.

### 3.2 Estimation hypothesis

Estimating a large number of free parameters from limited data poses a statistical challenge in the estimation of any model. We next quantified the influence of the number of free parameters and the number of observations on model stasis. We show that a high number of free parameters and a low number of observations can cause the model to become static.

In the simulated data, we found that both increasing the number of free parameters *DF* by including more parcels of the parcellation (i.e. increasing *N*) and decreasing the number of observations *O* by simulating fewer subjects increase model stasis. This is illustrated in **Figure 3**A where, compared to the models described above under *3*.*1*.*1* (plotted in the left panel), we increased the number of free parameters *DF* (middle panel), and additionally decreased the number of observations *O* (right panel). The area where the model becomes static (i.e. where mean maxFO is high, here shown in yellow) increases for both steps. In the full model, the standardised coefficients for the inverse of the number of free parameters *DF* is -0.16 (*p*=0.0001) and for the number of observations *O* is -0.21 (*p*<0.0001).

**Figure 3.**
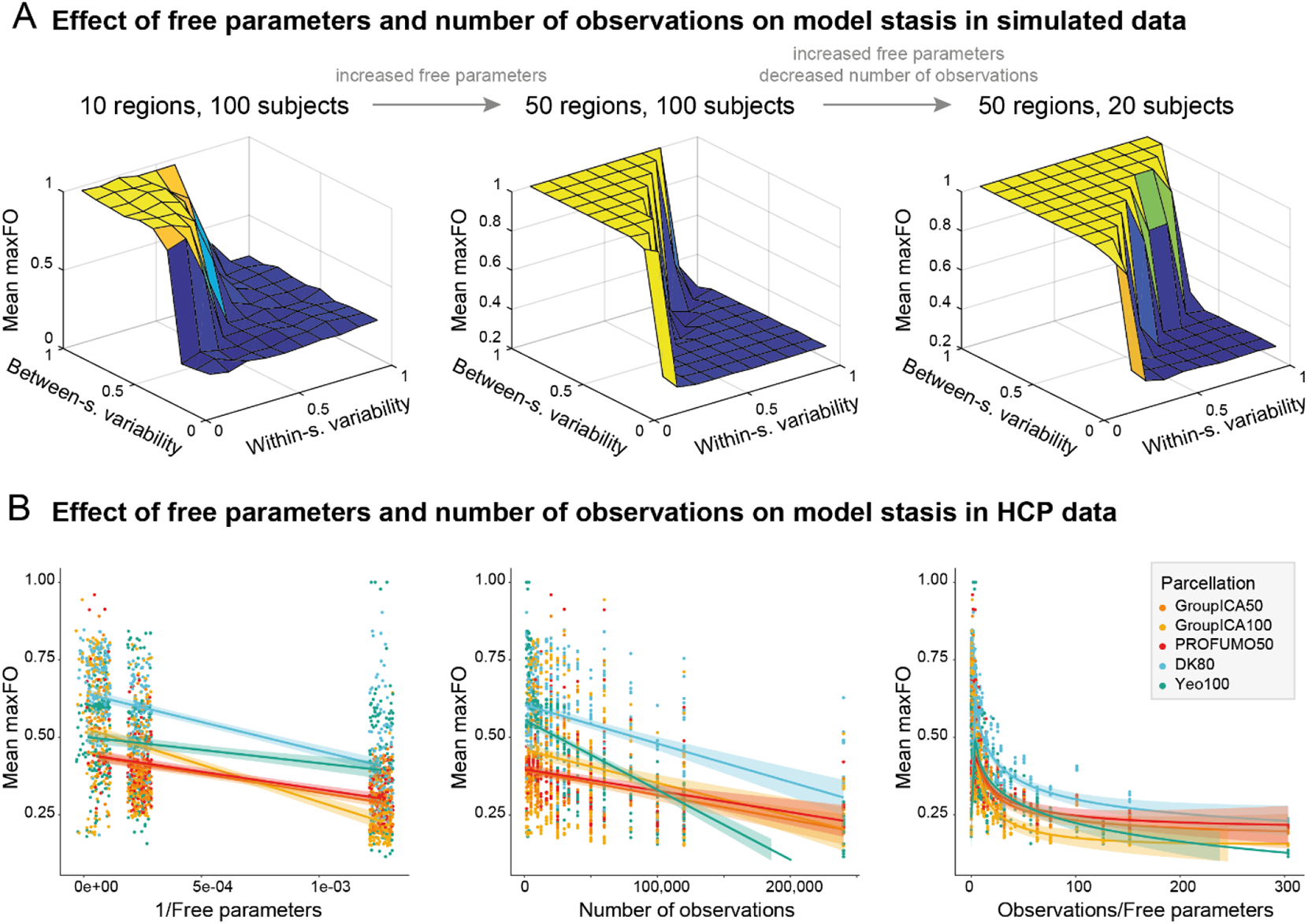
Evidence for the estimation hypothesis. **A)** In the simulated data, we first increased the number of free parameters DF by manipulating the number of parcels N included from the parcellation (middle panel). This increased the yellow area of the graph, i.e. the area where the time-varying FC model is static. In addition to increasing the number of free parameters DF, we then also decreased the number of observations O by simulating fewer subjects S (right panel). This further increased the area where the time-varying FC model is static, so that it is now only possible for the model to detect dynamics (blue area) when between-subject variability is very low and within-session variability is very high. **B)** In the real data, both decreasing the number of free parameters DF (left panel, where we show the inverse of the number of free parameters) and increasing the number of observations O (middle panel) reduce model stasis, as indicated by lower values in mean maxFO. Finally, the ratio of observations to free parameters (right panel) is a strong negative indicator of model stasis. This ratio is small in all models that are mostly static (high values of mean maxFO) and high in all models that are mostly dynamic (low values of mean maxFO). Given the differences between parcellations established in 3.1.2, we distinguish between parcellations in these plots. This distinction is here only for illustrative purposes and not included as random effects in the full SEM.

In the real data, the number of free parameters *DF* was manipulated by changing the number of parcels *N* as described under 2.1.2. As illustrated in **Figure 3**B, both decreasing the number of free parameters *DF* (left panel) and increasing the number of observations *O* (middle panel) decreased model stasis in the HCP data. A low ratio of number of observations *O* to free parameters *DF* (right panel) is a strong indicator of model stasis. Based on the finding that model stasis strongly depends on the parcellation, we here also plot these effects for each parcellation separately. Please note that we plot the inverse of the number of free parameters in the left panel, so that the values in the right panel are the product of the two previous plots. In the full SEM, the coefficient of the inverse of the number of free parameters *DF* is -0.50 (*p*<0.0001), the coefficient of the number of observations *O* is -0.30 (*p*<0.0001) and the coefficient of their interaction is -0.06 (*p*=0.02). As shown in the Supplementary Figures 1 and 2, reducing the number of free parameters *O* using the PCA- and HMM-PCA approaches similarly decreased model stasis. This effect was parcellation-dependent.

In a dataset with few observations, reducing the number of free parameters may therefore be vital for a time-varying FC model to detect dynamic changes in FC. If more data is available, it is possible to increase the number of free parameters and thus add detail to the model, e.g. by using a more fine-grained parcellation.

### 3.3 Synthesis of results

In order to compare the directed effect of all variables on model stasis, we finally modelled the influence of all factors described under 3.1 and **Error! Reference source not found**. using SEMs. We estimated separate models with a similar structure for the simulated and the real data, as described in 2.4. The structure and results of the SEM are summarised in **Figure 4**. The first part of each model uses FC similarity as the outcome measure, while the second part uses model stasis as the outcome. This structure allows differentiating for instance between a direct effect of the number of observations on model stasis and an indirect effect of the number of observations on model stasis via FC similarity.

**Figure 4.**
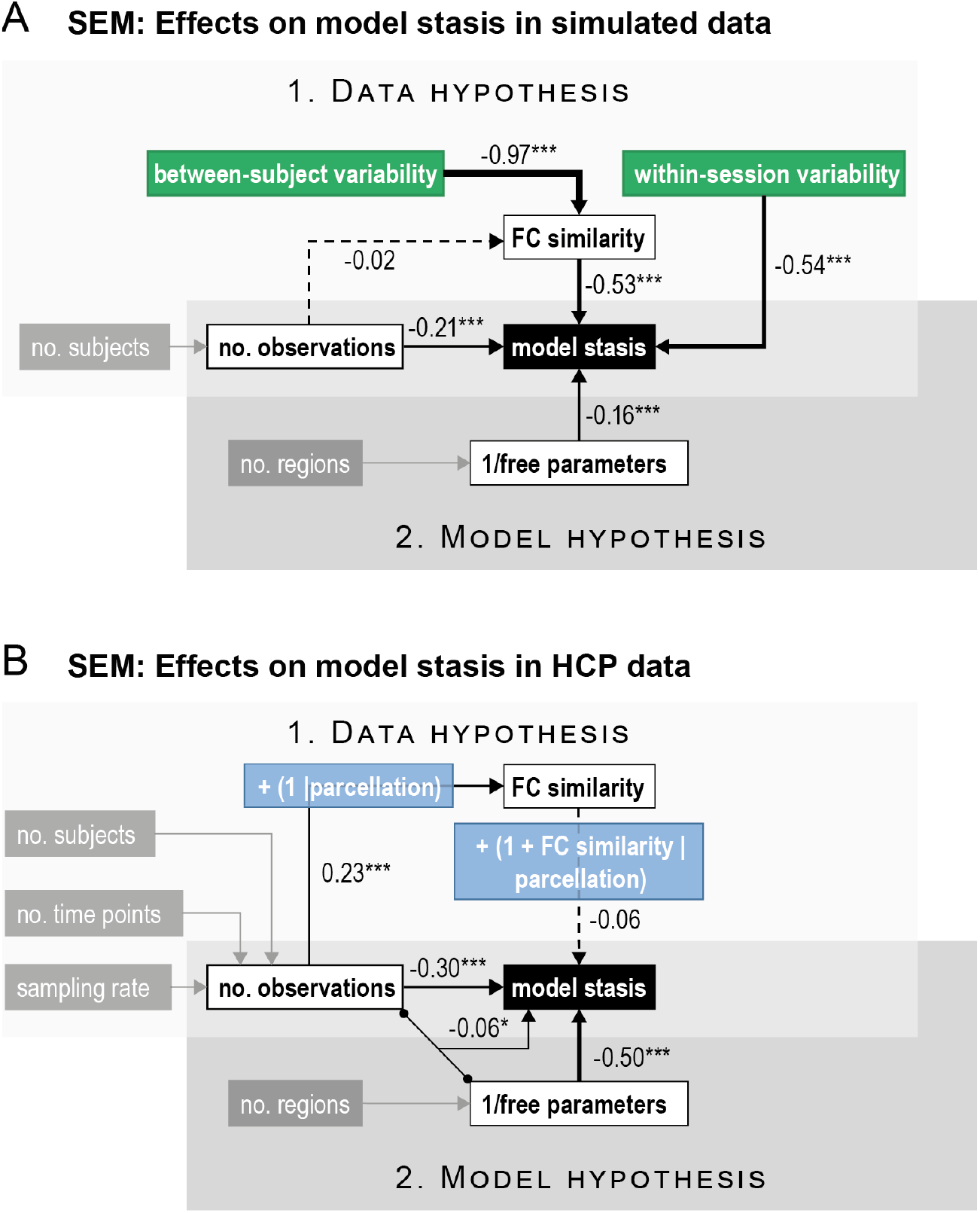
Full structural equation models (SEM). **A)** On simulated data, FC similarity (which is almost perfectly explained by between-subject variability) and within-session variability strongly affect model stasis, providing compelling evidence for the data hypothesis. Coefficients corresponding to the estimation hypothesis are smaller, but still both the number of free parameters and the number of observations significantly affect model stasis. **B)** On real data, the effect of the data hypothesis is less strong, as indicated by the smaller coefficients between FC similarity and model stasis. As explained in 3.1.2, variance in model stasis from the data hypothesis can be explained better by distinguishing between different parcellations. The number of free parameters and the number of observations, as well their interaction strongly affect model stasis. Here, grey boxes indicate variables that are not explicitly modelled in the SEM, but which are constituting parts of another variable. White boxes represent predictor variables. Green boxes are synthetically manipulated variables in the simulated data. Blue boxes specify random effects that affect the underlying link. The black box indicates the main outcome variable. Arrow thickness is scaled to the corresponding coefficient strength. Significance levels are indicated by asterisks: *** p < 0.001, ** p < 0.01, * p < 0.05.

The full models explain 95% and 86% variance in FC similarity, and 68% and 72% in model stasis, respectively, for simulated and real data. Comparing the two hypotheses, in the simulated data we found more evidence for the data hypothesis than for the estimation hypothesis. In the real data, however, the evidence supporting the estimation hypothesis dominates the data hypothesis, particularly the number of free parameters. An apparent difference between the simulated data and the real data is the effect of FC similarity (standardised coefficients of -0.53*** in simulated data and of -0.06 N.S. in real data). It is also important to note that we use only one parcellation in the simulations, but five different parcellations in the real data. We show above that, in the real data, 32% of variance in model stasis is explained by the random effects of parcellations. This indicates that, rather than using overall FC similarity as a single indicator of model stasis, it is important to distinguish between different parcellations. Another important difference between simulated and real data is that the amount of between-subject and within-session variability can only be directly manipulated on the synthetic data. However, between-subject and within-session variability often differ to a large extent between real datasets and are therefore an important consideration when applying time-varying FC models.

Overall, we found evidence for all hypothesised effects. At the level of the data hypothesis, we showed in the simulations that low between-subject and high within-session variability reduce model stasis. Additionally, on real data, we showed that the choice of parcellation strongly affects time-averaged FC, model stasis, and the relationship between them. At the level of the estimation hypothesis, we presented evidence that a larger number of observations and fewer free parameters reduce model stasis, both on simulated and on real data.

## 4 Discussion

The ability of a time-varying FC model to identify temporally changing states on fMRI data depends on numerous factors, and can be attributed both to aspects of the data and to aspects of the model. Our findings indicate that model stasis is affected by the actual variability in the data, the parcellation used to extract time courses, and the ratio of the number of available observations to the number of free parameters in the model. To summarise when these models can be satisfactorily applied, we have compiled a short list of practical recommendations in Conclusions.

We first showed that large differences between subjects and/or small within-session FC modulations can cause a time-varying FC model to become static. This can be explained by the data-driven, unsupervised nature of the model, which aims at describing the most salient features of a dataset without imposing specific constraints about the recurrence of states across subjects. We also showed that FC similarity, model stasis, and the relationship between them are affected by the parcellation. In the example parcellations we used here, the three data-driven parcellations on average resulted in lower model stasis (i.e. they were found to be better models from the specific point of view considered in this paper) than the examples of functional or anatomical parcellations. Although these conclusions might not necessarily generalise to all functional or anatomical parcellations, the effect was clear in this case. Understanding the reason behind these differences between parcellations is not straightforward, as there are several factors involved —such as differences in spatial distribution, cluster size, weighted vs. binary parcels, time course extraction, etc.— that may contribute to these differences, and these have not been explicitly tested here. Assuming the presence of “true” functional clusters in the data, data-driven functional parcellations are more likely to detect these clusters as parcels, resulting in a more efficient estimation of the temporal variance of these clusters. In theory, anatomical and *a priori* functional parcellations may, for example, split “true” functional clusters into several parcels, which could affect the balance between between-subject and within-session variability in an artefactual manner.

Second, we showed that a high number of free parameters in the model can cause the model to become static, especially if too few observations are available to estimate these parameters. Here we showed that the model may become static when too many free parameters need to be estimated. This is because, if the data available for the estimation of time-varying FC is insufficient, avoiding all state switches might be the most parsimonious solution in terms of the model inference. This implies that the estimation of time-varying FC is a trade-off between the level of detail in the spatial domain and the accuracy of the temporal estimation.

Importantly, the factors we considered here are not exhaustive and therefore other variables related to overall data quality and model characteristics might also be relevant. In particular, a large dissonance between the model specification and the realities of the data could also be a reason why we could not detect time-varying FC. For instance, if temporal modulations in first-order statistics (the average pattern of activity within a state —i.e. the mean of the Gaussian distribution) were temporally independent from modulations in time-varying FC, this would violate the assumptions of the HMM and could potentially affect model stasis; in this case, modelling the mean as a separate temporal process would likely improve the estimation of time-varying FC (Pervaiz *et al*., 2021).

It also remains to be seen how exactly model stasis may occur in other kinds of models (such as a mixture of Gaussian distributions (Bishop, 2006)), although the logic of our conclusions is likely to remain valid. It should also be noted that we here only focussed on model stasis, because it is among the most fundamental measures of performance of a time-varying FC model. However, other evaluative measures, such as the ability to predict individual traits and behaviour may be of interest when evaluating time-varying FC model performance, as shown in Pervaiz *et al*. (2021); Pervaiz *et al*. (2020); Vidaurre *et al*. (2021) and many other works. It is likely that some of the variables we here showed to reduce model stasis, such as higher similarity between subjects and fewer free parameters (as obtained, e.g., through a coarser parcellation), would indeed be disadvantageous when considering other evaluative measures or when conducting a time-averaged FC study. The assessment of this will be the object of future work.

## 5 Conclusion

As we outlined in this article, the ability to estimate time-varying FC in fMRI data depends on several factors, which should be considered when planning and conducting a time-varying FC study. To avoid a time-varying FC model becoming static, we provide the following recommendations:

1. *Preprocessing*: Special care should be taken in reducing artefactual between-subject differences, e.g. by optimising registration and removing subject-specific artefacts, and in preserving temporal variance by refraining from preprocessing steps that average over time points like motion scrubbing or other more “aggressive” clean-up strategies. Testing similarity in time-averaged FC between subjects may in some cases be useful as an indicator of problematic between-subject variability, but it can also be misleading in certain parcellations.
2. *Time course extraction*: The choice of parcellation used to extract time courses should be considered when planning a time-varying FC study. The data-driven functional parcellations we used here, such as Group ICA approaches, perform better than the examples we used for functional or anatomical parcellations in detecting temporal changes in FC.
3. *Model complexity*: The number of free parameters should ideally be not too large in relation to the number of observations, e.g. by using a parcellation with fewer parcels or components if necessary. Other options to reduce the number of free parameters include dimensionality reduction, e.g. using Principal Component Analysis (PCA), which however may affect the model in other ways (Vidaurre, 2021). Based on the HCP-dataset, we estimate as a rule of thumb that the ratio of observations to free parameters should not be inferior to 10.

In summary, meeting these requirements may help improving the robustness and reliability of time-varying FC methods and eventually increase replicability (Choe *et al*., 2017).

## Data and code availability statement

The real dataset used in this study has been made publicly available by the WU-Minn Consortium in ConnectomeDB https://db.humanconnectome.org/. Code to reproduce the simulated dataset and to replicate the analyses of this study is available under https://github.com/ahrends/mixing. The HMM-MAR toolbox used to run the HMMs is available under https://github.com/OHBA-analysis/HMM-MAR.

## Acknowledgments

We would like to thank Stephen Smith for helpful discussions. Real data were provided by the Human Connectome Project, WU-Minn Consortium (Principal Investigators: David Van Essen and Kamil Ugurbil; 1U54MH091657) funded by the 16 NIH Institutes and Centers that support the NIH Blueprint for Neuroscience Research; and by the McDonnell Center for Systems Neuroscience at Washington University. DV is supported by a Novo Nordisk Foundation Emerging Investigator Fellowship (NNF19OC-0054895) and an ERC Starting Grant (ERC-StG-2019-850404). CA is funded by the Danish National Research Foundation (DNRF117). UP is funded by an MRC Mental Health Data Path Finder award (PI Clare Mackay) MC/PC/17215. The Wellcome Centre for Integrative Neuroimaging is supported by core funding from the Wellcome Trust (203139/Z/16/Z). MWW’s research is supported by the NIHR Oxford Health Biomedical Research Centre, the Wellcome Trust (106183/Z/14/Z, 215573/Z/19/Z), the New Therapeutics in Alzheimer’s Diseases (NTAD) study supported by UK MRC and the Dementia Platform UK (RG94383/RG89702) and the EU-project euSNN (MSCA-ITN H2020-860563).

## References

Battaglia, D., Boudou, T., Hansen, E.C.A., Lombardo, D., Chettouf, S., Daffertshofer, A., McIntosh, A.R., Zimmermann, J., Ritter, P., Jirsa, V., 2020. Dynamic Functional Connectivity between order and randomness and its evolution across the human adult lifespan. NeuroImage 222, 117156. doi:10.1016/j.neuroimage.2020.117156.

Beckmann, C.F., 2012. Modelling with independent components. NeuroImage 62, 891–901. doi:10.1016/j.neuroimage.2012.02.020.

Beckmann, C.F., Mackay, C.E., Filippini, N., Smith, S.M., 2009. Group comparison of resting-state FMRI data using multi-subject ICA and dual regression. NeuroImage 47, S148. doi:10.1016/S1053-8119(09)71511-3.

Bijsterbosch, J.D., Woolrich, M.W., Glasser, M.F., Robinson, E.C., Beckmann, C.F., Van Essen, D.C., Harrison, S.J., Smith, S.M., 2018. The relationship between spatial configuration and functional connectivity of brain regions. eLife 7, e32992. doi:10.7554/eLife.32992.

Bishop, C., 2006. Pattern Recognition and Machine Learning. Springer-Verlag New York.

Breakspear, M., 2017. Dynamic models of large-scale brain activity. Nat Neurosci 20, 340–352. doi:10.1038/nn.4497.

Cabral, J., Vidaurre, D., Marques, P., Magalhaes, R., Silva Moreira, P., Miguel Soares, J., Deco, G., Sousa, N., Kringelbach, M.L., 2017. Cognitive performance in healthy older adults relates to spontaneous switching between states of functional connectivity during rest. Sci Rep 7, 5135. doi:10.1038/s41598-017-05425-7.

Calhoun, Vince D., Miller, R., Pearlson, G., Adalı, T., 2014. The Chronnectome: Time-Varying Connectivity Networks as the Next Frontier in fMRI Data Discovery. Neuron 84, 262–274. doi:10.1016/j.neuron.2014.10.015.

Casorso, J., Kong, X., Chi, W., Van De Ville, D., Yeo, B.T.T., Liégeois, R., 2019. Dynamic mode decomposition of resting-state and task fMRI. NeuroImage 194, 42–54. doi:10.1016/j.neuroimage.2019.03.019.

Choe, A.S., Nebel, M.B., Barber, A.D., Cohen, J.R., Xu, Y., Pekar, J.J., Caffo, B., Lindquist, M.A., 2017. Comparing test-retest reliability of dynamic functional connectivity methods. NeuroImage 158, 155–175. doi:10.1016/j.neuroimage.2017.07.005.

Dai, M., Zhang, Z., Srivastava, A., 2019. Discovering common change-point patterns in functional connectivity across subjects. Medical Image Analysis 58. doi:10.1016/j.media.2019.101532.

Deco, G., Vidaurre, D., Kringelbach, M.L., 2021. Revisiting the global workspace orchestrating the hierarchical organization of the human brain. Nature Human Behaviour 5, 497–511. doi:10.1038/s41562-020-01003-6.

Desikan, R.S., Ségonne, F., Fischl, B., Quinn, B.T., Dickerson, B.C., Blacker, D., Buckner, R.L., Dale, A.M., Maguire, R.P., Hyman, B.T., Albert, M.S., Killiany, R.J., 2006. An automated labeling system for subdividing the human cerebral cortex on MRI scans into gyral based regions of interest. Neuroimage 31, 968–980. doi:10.1016/j.neuroimage.2006.01.021.

Eickhoff, S.B., Yeo, B.T.T., Genon, S., 2018. Imaging-based parcellations of the human brain. Nature Reviews Neuroscience 19, 672–686. doi:10.1038/s41583-018-0071-7.

Fornito, A., Bullmore, E.T., 2010. What can spontaneous fluctuations of the blood oxygenationlevel-dependent signal tell us about psychiatric disorders? Current Opinion in Psychiatry 23, 239–249. doi:10.1097/YCO.0b013e328337d78d.

Fries, P., 2005. A mechanism for cognitive dynamics: neuronal communication through neuronal coherence. Trends Cogn Sci 9, 474–480. doi:10.1016/j.tics.2005.08.011.

Glasser, M.F., Sotiropoulos, S.N., Wilson, J.A., Coalson, T.S., Fischl, B., Andersson, J.L., Xu, J., Jbabdi, S., Webster, M., Polimeni, J.R., Van Essen, D.C., Jenkinson, M., 2013. The minimal preprocessing pipelines for the Human Connectome Project. Neuroimage 80, 105–124. doi:10.1016/j.neuroimage.2013.04.127.

Gonzalez-Castillo, J., Bandettini, P.A., 2018. Task-based dynamic functional connectivity: Recent findings and open questions. Neuroimage 180, 526–533. doi:10.1016/j.neuroimage.2017.08.006.

Griffanti, L., Salimi-Khorshidi, G., Beckmann, C.F., Auerbach, E.J., Douaud, G., Sexton, C.E., Zsoldos, E., Ebmeier, K.P., Filippini, N., Mackay, C.E., Moeller, S., Xu, J., Yacoub, E., Baselli, G., Ugurbil, K., Miller, K.L., Smith, S.M., 2014. ICA-based artefact removal and accelerated fMRI acquisition for improved resting state network imaging. Neuroimage 95, 232–247. doi:10.1016/j.neuroimage.2014.03.034.

Harrison, S.J., Woolrich, M.W., Robinson, E.C., Glasser, M.F., Beckmann, C.F., Jenkinson, M., Smith, S.M., 2015. Large-scale Probabilistic Functional Modes from resting state fMRI. NeuroImage 109, 217–231. doi:10.1016/j.neuroimage.2015.01.013.

Hindriks, R., Adhikari, M.H., Murayama, Y., Ganzetti, M., Mantini, D., Logothetis, N.K., Deco, G., 2016. Can sliding-window correlations reveal dynamic functional connectivity in resting-state fMRI? Neuroimage 127, 242–256. doi:10.1016/j.neuroimage.2015.11.055.

Lefcheck, J.S., 2016. piecewiseSEM: Piecewise structural equation modelling in r for ecology, evolution, and systematics. Methods in Ecology and Evolution 7, 573–579. doi:10.1111/2041-210X.12512.

Lehmann, B.C.L., White, S.R., Henson, R.N., Cam, C.A.N., Geerligs, L., 2017. Assessing dynamic functional connectivity in heterogeneous samples. NeuroImage 157, 635–647. doi:10.1016/j.neuroimage.2017.05.065.

Liégeois, R., Laumann, T.O., Snyder, A.Z., Zhou, J., Yeo, B.T.T., 2017. Interpreting temporal fluctuations in resting-state functional connectivity MRI. NeuroImage 163, 437–455. doi:10.1016/j.neuroimage.2017.09.012.

Liegeois, R., Li, J., Kong, R., Orban, C., Van De Ville, D., Ge, T., Sabuncu, M.R., Yeo, B.T.T., 2019. Resting brain dynamics at different timescales capture distinct aspects of human behavior. Nat Commun 10, 2317. doi:10.1038/s41467-019-10317-7.

Lindquist, M.A., Xu, Y., Nebel, M.B., Caffo, B.S., 2014. Evaluating dynamic bivariate correlations in resting-state fMRI: A comparison study and a new approach. NeuroImage 101, 531–546. doi:10.1016/j.neuroimage.2014.06.052.

Lurie, D.J., Kessler, D., Bassett, D.S., Betzel, R.F., Breakspear, M., Kheilholz, S., Kucyi, A., Liégeois, R., Lindquist, M.A., McIntosh, A.R., Poldrack, R.A., Shine, J.M., Thompson, W.H., Bielczyk, N.Z., Douw, L., Kraft, D., Miller, R.L., Muthuraman, M., Pasquini, L., Razi, A., Vidaurre, D., Xie, H., Calhoun, V.D., 2019. Questions and controversies in the study of time-varying functional connectivity in resting fMRI. Network Neuroscience 4, 30–69. doi:10.1162/netn_a_00116.

MATLAB, 2016. R2016b. The MathWorks Inc.: Natick, Massachusetts.

Pervaiz, U., Vidaurre, D., Gohil, C., Smith, S.M., Woolrich, M.W., 2021. Multi-dynamic Modelling Reveals Strongly Time-varying Resting fMRI Correlations. bioRxiv, 2021.2006.2023.449584. doi:10.1101/2021.06.23.449584.

Pervaiz, U., Vidaurre, D., Woolrich, M.W., Smith, S.M., 2020. Optimising network modelling methods for fMRI. NeuroImage 211, 116604. doi:https://doi.org/10.1016/j.neuroimage.2020.116604.

Popovych, O.V., Jung, K., Manos, T., Diaz-Pier, S., Hoffstaedter, F., Schreiber, J., Yeo, B.T.T., Eickhoff, S.B., 2021. Inter-subject and inter-parcellation variability of resting-state whole-brain dynamical modeling. Neuroimage, 118201. doi:10.1016/j.neuroimage.2021.118201.

Quinn, A.J., Vidaurre, D., Abeysuriya, R., Becker, R., Nobre, A.C., Woolrich, M.W., 2018. Task-Evoked Dynamic Network Analysis Through Hidden Markov Modeling. Frontiers in Neuroscience 12. doi:10.3389/fnins.2018.00603.

Riccelli, R., Passamonti, L., Duggento, A., Guerrisi, M., Indovina, I., Toschi, N. (2017) Dynamic internetwork connectivity in the human brain. In: Proceedings of the Annual International Conference of the IEEE Engineering in Medicine and Biology Society, EMBS. pp. 3313–3316.

Salimi-Khorshidi, G., Douaud, G., Beckmann, C.F., Glasser, M.F., Griffanti, L., Smith, S.M., 2014. Automatic denoising of functional MRI data: combining independent component analysis and hierarchical fusion of classifiers. Neuroimage 90, 449–468. doi:10.1016/j.neuroimage.2013.11.046.

Schaefer, A., Kong, R., Gordon, E.M., Laumann, T.O., Zuo, X.N., Holmes, A.J., Eickhoff, S.B., Yeo, B.T.T., 2018. Local-Global Parcellation of the Human Cerebral Cortex from Intrinsic Functional Connectivity MRI. Cereb Cortex 28, 3095–3114. doi:10.1093/cercor/bhx179.

Shipley, B., 2000. A New Inferential Test for Path Models Based on Directed Acyclic Graphs. Structural Equation Modeling: A Multidisciplinary Journal 7, 206–218. doi:10.1207/S15328007SEM0702_4.

Smith, S.M., Beckmann, C.F., Andersson, J., Auerbach, E.J., Bijsterbosch, J., Douaud, G., Duff, E., Feinberg, D.A., Griffanti, L., Harms, M.P., Kelly, M., Laumann, T., Miller, K.L., Moeller, S., Petersen, S., Power, J., Salimi-Khorshidi, G., Snyder, A.Z., Vu, A.T., Woolrich, M.W., Xu, J., Yacoub, E., Uğurbil, K., Van Essen, D.C., Glasser, M.F., 2013a. Resting-state fMRI in the Human Connectome Project. Neuroimage 80, 144–168. doi:10.1016/j.neuroimage.2013.05.039.

Smith, S.M., Beckmann, C.F., Andersson, J., Auerbach, E.J., Bijsterbosch, J., Douaud, G., Duff, E., Feinberg, D.A., Griffanti, L., Harms, M.P., Kelly, M., Laumann, T., Miller, K.L., Moeller, S., Petersen, S., Power, J., Salimi-Khorshidi, G., Snyder, A.Z., Vu, A.T., Woolrich, M.W., Xu, J., Yacoub, E., Uğurbil, K., Van Essen, D.C., Glasser, M.F., Consortium, W.U.-M.H., 2013b. Resting-state fMRI in the Human Connectome Project. NeuroImage 80, 144–168. doi:10.1016/j.neuroimage.2013.05.039.

Smith, S.M., Vidaurre, D., Beckmann, C.F., Glasser, M.F., Jenkinson, M., Miller, K.L., Nichols, T.E., Robinson, E.C., Salimi-Khorshidi, G., Woolrich, M.W., Barch, D.M., Uğurbil, K., Van Essen, D.C., 2013c. Functional connectomics from resting-state fMRI. Trends in Cognitive Sciences 17, 666–682. doi:10.1016/j.tics.2013.09.016.

Sporns, O., Faskowitz, J., Teixeira, A.S., Cutts, S.A., Betzel, R.F., 2021. Dynamic expression of brain functional systems disclosed by fine-scale analysis of edge time series. Network Neuroscience 5, 405–433. doi:10.1162/netn_a_00182.

Stevner, A.B.A., Vidaurre, D., Cabral, J., Rapuano, K., Nielsen, S.F.V., Tagliazucchi, E., Laufs, H., Vuust, P., Deco, G., Woolrich, M.W., Van Someren, E., Kringelbach, M.L., 2019. Discovery of key whole-brain transitions and dynamics during human wakefulness and non-REM sleep. Nat Commun 10, 1035. doi:10.1038/s41467-019-08934-3.

Team, R.C. (2020) R: A language and environment for statistical computing. R Foundation for Statistical Computing: Vienna, Austria.

Van Essen, D.C., Smith, S.M., Barch, D.M., Behrens, T.E., Yacoub, E., Ugurbil, K., 2013. The WU-Minn Human Connectome Project: an overview. Neuroimage 80, 62–79. doi:10.1016/j.neuroimage.2013.05.041.

Van Essen, D.C., Ugurbil, K., Auerbach, E., Barch, D., Behrens, T.E.J., Bucholz, R., Chang, A., Chen, L., Corbetta, M., Curtiss, S.W., Della Penna, S., Feinberg, D., Glasser, M.F., Harel, N., Heath, A.C., Larson-Prior, L., Marcus, D., Michalareas, G., Moeller, S., Oostenveld, R., Petersen, S.E., Prior, F., Schlaggar, B.L., Smith, S.M., Snyder, A.Z., Xu, J., Yacoub, E., Consortium, W.U.-M.H., 2012. The Human Connectome Project: a data acquisition perspective. NeuroImage 62, 2222–2231. doi:10.1016/j.neuroimage.2012.02.018.

Vidaurre, D., 2021. A new model for simultaneous dimensionality reduction and time-varying functional connectivity estimation. PLOS Computational Biology 17, e1008580. doi:10.1371/journal.pcbi.1008580.

Vidaurre, D., Llera, A., Smith, S.M., Woolrich, M.W., 2021. Behavioural relevance of spontaneous, transient brain network interactions in fMRI. NeuroImage 229, 117713. doi:10.1016/j.neuroimage.2020.117713.

Vidaurre, D., Quinn, A.J., Baker, A.P., Dupret, D., Tejero-Cantero, A., Woolrich, M.W., 2016. Spectrally resolved fast transient brain states in electrophysiological data. Neuroimage 126, 81–95. doi:10.1016/j.neuroimage.2015.11.047.

Vidaurre, D., Smith, S.M., Woolrich, M.W., 2017. Brain network dynamics are hierarchically organized in time. Proc Natl Acad Sci U S A 114, 12827–12832. doi:10.1073/pnas.1705120114.

Voytek, B., Knight, R.T., 2015. Dynamic network communication as a unifying neural basis for cognition, development, aging, and disease. Biol Psychiatry 77, 1089–1097. doi:10.1016/j.biopsych.2015.04.016.

Xie, H., Calhoun, V.D., Gonzalez-Castillo, J., Damaraju, E., Miller, R., Bandettini, P.A., Mitra, S., 2018. Whole-brain connectivity dynamics reflect both task-specific and individual-specific modulation: A multitask study. Neuroimage 180, 495–504. doi:10.1016/j.neuroimage.2017.05.050.

Zalesky, A., Fornito, A., Cocchi, L., Gollo, L.L., Breakspear, M., 2014. Time-resolved resting-state brain networks. Proceedings of the National Academy of Sciences 111, 10341–10346. doi:10.1073/pnas.1400181111.

Zamani Esfahlani, F., Jo, Y., Faskowitz, J., Byrge, L., Kennedy, D.P., Sporns, O., Betzel, R.F., 2020. High-amplitude cofluctuations in cortical activity drive functional connectivity. Proceedings of the National Academy of Sciences 117, 28393–28401. doi:10.1073/pnas.2005531117.

